# Effect of tuft-IL-25-ILC2 pathway on mouse small intestine stimulated by *Trichinella spiralis* ES antigen

**DOI:** 10.1101/2023.02.23.529658

**Authors:** Bai Jie, HE Ling, Napisha Jureti

## Abstract

**Objective:** To explore the regulatory effect of *Trichinella spiralis* ES antigen on the intestinal immune function, we observed the immune response of mouse small intestine to this antigen by quantifying the changes in the related cytokines of Tuft-IL-25-ILC2 pathway.

**Methods:** A total of 30 BALB/c female mice were randomly divided into three groups: control group, *Trichinella* excretion-secretion (ES) antigen stimulation group, and IL-25 blocking group. The mice in the control group were injected intraperitoneally with PBS; those in the ES antigen stimulation group were injected intraperitoneally with ES antigen once per day for 7 consecutive days; and those in the IL-25 blocking group were injected intraperitoneally, first with anti-mouse IL-25 monoclonal antibody and 3 days later with ES antigen. Alcian blue-nuclear fast red staining was employed to observe the changes in the number of small intestinal goblet cells. The number of Tuft cells was determined by immunofluorescence chemical analysis, and the expression levels of IL-25, IL-13, IL-25R, Pou2f3, and RORα mRNA were quantified by RT-PCR.

**Key results:** The results of Alcian blue-nuclear fast red staining showed that, compared with the control group, the number of goblet cells in the small intestinal tissue of mice in the ES antigen stimulation group increased, and the difference was statistically significant (P<0.05). Compared with the ES antigen stimulation group, the number of goblet cells in the small intestinal tissue of the mice in the IL-25 blocking group was slightly decreased (P<0.05). Immunofluorescence analysis showed that, compared with the control group, the number of Tuft cells in the ES antigen-stimulated group increased (P<0.05), while in the IL-25 blocking group, the number of Tuft cells was decreased compared with that in the ES antigen stimulation group (P<0.05). RT-PCR analysis showed that, compared with the control group, the mRNA expression levels of IL-25, IL-13, IL- 25R, ROR*α* and Pou2f3 in the small intestine of mice in the ES antigen stimulation group were increased (P<0.05); compared with the ES antigen stimulation group, the IL-25, IL-13, IL-25R, ROR*α* and Pou2f3 mRNA expressions in the tissues of the mice in the IL-25 blocking group were decreased, and the difference was statistically significant (P<0.05).

**Conclusion:** The small intestinal mucosa of mice stimulated by *Trichinella* ES antigen may possess an immune function via the Tuft-IL-25-ILC2 pathway.

## INTRODUCTION

*Trichinella spiralis* is the largest intracellular parasitic nematode, which causes trichinosis, a zoonotic disease [1]. The clinical symptoms are complex and diverse, including nausea, vomiting, fever, diarrhea, etc., and the pathogenic diagnosis is difficult. In the process of host infection, the parasite can produce various antigens. Among them, the excretory-secretory (ES) antigen is directly exposed to the host immune system, which is the main target antigen for inducing host immune response [2]. It can regulate autoimmunity and maintain immune system homeostasis [3]. Studies have shown that *Trichinella spiralis* exerts an immunomodulatory effect on the host immune response through its excretion and secretion products, resulting in the polarization of the host immune response to the Th2 phenotype [4]. Tuft cells, as a small proportion of intestinal epithelial cells, play an important role in recognizing and responding to parasitic infections [5]. Studies have shown that tuft cells can receive parasite infection signals and release the cytokine IL-25 to trigger a type II innate immune response, which ultimately promotes the expulsion of parasites from the body and is detrimental to parasitic infection.

In our experiment, the ES antigen of *Trichinella* muscle larvae was used to stimulate the small intestine of healthy mice, and the changes in related cytokines of the Tuft-IL-25-ILC2 pathway in the small intestine of mice were observed. The expression of mouse intestinal goblet cells was determined by Alcian blue-nuclear fast red staining, the alterations in the number of Tuft cells by immunofluorescence chemical analysis were evaluated, and RT-PCR was used to detect the expression levels of IL-25, IL-13, IL-25R, RORα, and Pou2f3 mRNA in the small intestinal tissue. Our aims were to explore the detailed mechanism of action between *Trichinella* and the immune system of the Tuft-IL-25-ILC2 pathway in the small intestine of the host, and to provide new ideas for the control of clinical trichinosis infections.

## MATERIALS AND METHODS

### 1.1 Experimental animals

The *Trichinella spiralis* specimens used in this study were obtained from the Heilongjiang Provincial Key Laboratory of Zoonotic Diseases of Animal Origin, and were kept in Kunming mice. A total of 30 SPF grade 6-8 week old female BALB/c female mice were used, weighing 18- 22 g each. All animals were provided by the Animal Experimental Center of Xinjiang Medical University, production license number: SCXK (new) 2018-0002.

### 1.2 Main reagents and instruments

The bicinchoninic acid (BCA) protein quantification kit was obtained from Solebao Company, and the RNA rapid extraction kit was purchased from Biomed Biotechnology Co., Ltd. Microtome (Leica, Germany). Other equipment included an embedding machine (Changzhou Zhongwei Electronic Instrument Co., Ltd.), tissue dehydrator (Wuhan Tianzhirui Medical Technology Co., Ltd.), and slicer (Shanghai Precision Instrument Co., Ltd.).

### 1.3 Preparation of *Trichinella* ES antigens [5]

A total of 400 Kunming mice were orally infected with muscle larvae obtained from *Trichinella* mice preserved in our laboratory. After 45 days, the mice were slaughtered. *Trichinella* muscle larvae were obtained by artificial gastric juice digestion and collected by Bellman’s device. The collected worms were washed 5 times with sterile saline containing double antibodies, added to 1640 medium at a density of 5000 cells/mL, and incubated at 37°C and 5% CO_2_ for 48 h. The culture medium was collected and centrifuged. The supernatant was put into a dialysis bag for dialysis, and then concentrated with PEG 20000. The obtained ES antigen was filtered through a 0.22 μm membrane and detected by SDS-PAGE. The protein content was determined, and samples were stored at −20°C for future use.

### 1.4 Grouping and sampling of animal experiments

A total of 30 BALB/c female mice were randomly divided into 3 groups: control group, *Trichinella* excretion-secretion (ES) antigen stimulation group and IL-25 blocking group. The mice in the control group were intraperitoneally injected with PBS. ES antigen-stimulated mice were intraperitoneally injected with *Trichinella* sp. ES antigen. Meanwhile, mice in the IL-25 blocking group were injected intraperitoneally with anti-mouse IL-25 monoclonal antibody, and then with *Trichinella* sp. ES antigen three days later, once a day for 7 consecutive days. The mice in each group were sacrificed after 12 hours of fasting (free access to water) after the last injection. The abdomen was incised along the midline, the small intestine of the mice was exposed and washed; one part was fixed with 4% paraformaldehyde, and the other part was frozen at −80°C.

### 1.5 Alcian blue-nuclear fast red staining [6] and immunofluorescence observation

The small intestine was separated, specimens were collected, and about 1 cm of small intestine tissue was taken. After fixation with 4% paraformaldehyde, routine dehydration, and embedding sections, Alcian blue-nuclear fast red staining and immunofluorescence experiments were performed to observe the changes in small intestine tissue under a microscope.

### 1.6 RT-PCR detection of IL-25, IL-13, IL-25R, Pou2f3, and RORα mRNA expression levels in the mouse small intestine

First, 30 mg of colon tissue was weighed, total RNA was extracted in a clean bench according to the method of RNA rapid extraction kit, and the content and purity of total RNA was determined with a nucleic acid quantifier. cDNA was synthesized according to the kit method and stored at - 20°C. The gene sequence was searched according to the NCBI Gene bank and the product was verified with NCBI Primer BLAST function. The primer screening conditions were as follows: PCR product length: 100-200 bp; annealing temperature: 60°C, and the selected primer sequences were synthesized by Shanghai Sangong Biotechnology Co., Ltd. The primer information table is shown in Table 1. The gene expression was calculated according to the 2^-ΔΔCT^ method, and the standard curve and amplification efficiency of the gene were obtained. Each sample consisted of 2 duplicate wells, the total reaction system was 20 μL, the reaction system was 10 μL of TB Green Premix Ex Taq II, 0.8 μL of upstream and downstream primers, 6 μL of sterile water, 0.4 μL of ROX, and 2 μL of cDNA. The reaction conditions were: 95°C for 30 s, 95°C for 5 s, 60°C for 34 s, 95°C for 15 s, 60°C for 1 h, and 95°C for 15 s. Correlation calculation was performed according to the CT value of the sample, and whether the PCR product was specific was judged according to the melting curve.

**Table 1.**
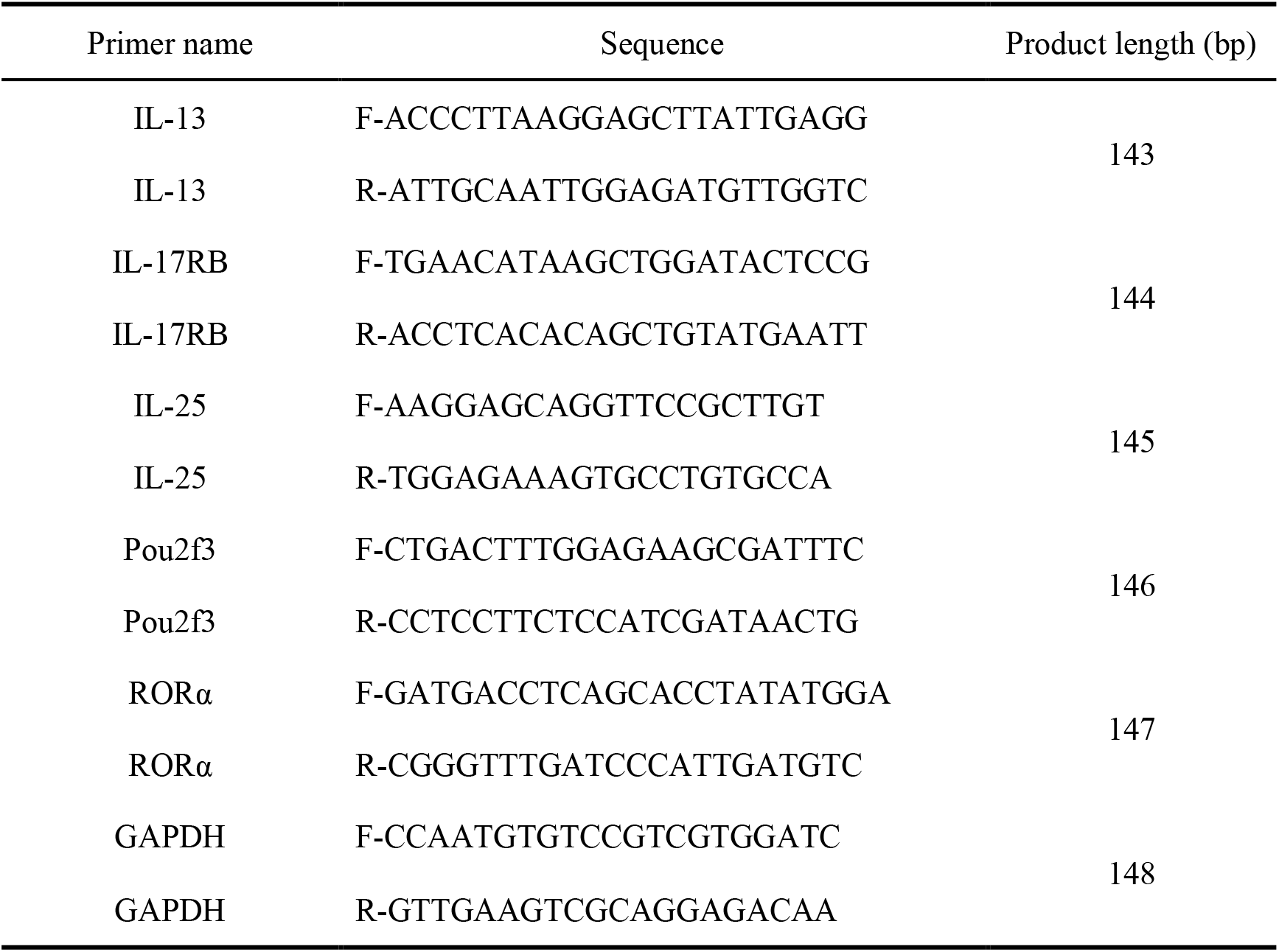
Primer sequence information table.

#### Statistical analysis

GraphPad Prism7 software was used for drawing, and SPSS23.0 software was employed for data analysis. All measurement data were expressed as mean ± standard deviation 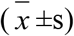. Oneway analysis of variance (ANOVA) was used to compare samples among multiple groups, and the LSD method was utilized for pairwise comparison when the variance was homogeneous. *P*<0.05 indicated a statistically significant difference.

## RESULTS

### 2.1 Preparation of *Trichinella* ES antigen

The collected muscle larvae were cultured in 1640 medium to obtain the *Trichinella spiralis* ES antigen. After the obtained culture medium was dialyzed, concentrated, filtered and the protein content was determined, the preparation and collection of ES antigen were detected by SDS-PAGE. The results showed that the target protein bands were mainly located at 35-53 kD (Figure 1), indicating that the prepared ES antigens were of good quality.

### 2.2 Changes in intestinal goblet cells stimulated by *Trichinella* ES antigens

The small intestine of mice was observed under the microscope of Alcian blue staining. Compared with the control group, the ES antigen stimulation group had more acidic mucus, the number of goblet cells increased, and the cell volume became larger; compared with the ES antigen stimulation group, the number of goblet cells in the IL-25 blocking group was relatively decreased, and the difference was statistically significant (Figure 2 and Figure 3).

### 2.3 Changes in the number of small intestinal Tuft cells stimulated by *Trichinella* ES antigen

The number of Tuft cells in the ES antigen stimulation group increased (*P*<0.05) compared with the control group, while the number of Tuft cells in the IL-25 blocking group decreased (*P*<0.05) compared with the ES antigen stimulation group (Figure 4 and Figure 5).

### 2.4 RT-PCR detection of cytokine mRNA expression levels in mouse small intestine

We further detected the mRNA expression of cytokines related to the IL-25-ILC2 pathway in the small intestine of mice by RT-PCR experiments (Table 2). Through experimental observation, it was found that, compared with the control group, the expression levels of IL-13, IL-25, RORα, Pou2f3, and IL-25R mRNA in the small intestinal tissue of the mice in the *Trichinella* ES antigen stimulation group were increased (*P*<0.05). Meanwhile, compared with the ES antigen stimulation group, the mRNA expression levels of IL-13, IL-25, RORα, Pou2f3, and IL-25R in the small intestinal tissue of the mice in the IL-25 blocking group were decreased, and the difference was statistically significant (*P*<0.05). (Figure 6)

**Table2.** The mRNA expression levels of small intestinal tissue-related cytokines in mice 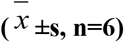.

| Study group | IL-13 | IL-25 | IL-25R | pou2f3 | RORα |
| --- | --- | --- | --- | --- | --- |
| control group | 1.00±0.14 | 1.00±0.16 | 1.00±0.15 | 1.00±0.14 | 1.00±0.14 |
| ES antigen stimulation group | 1.70±0.14* | 1.41±0.13* | 1.98±0.48* | 1.54±0.28* | 1.31±0.05* |
| IL-25 blocking group | 1.12±1.19# | 1.18±0.04# | 1.21±0.29# | 1.04±0.06# | 1.13±0.05# |

Note: Compared with the control group, \* $P<0.05$ ; compared with the ES antigen stimulation group, # $P<0.05$ .

## 3 Discussion

*Trichinella* can interact with the host immune system through ES antigens at different developmental stages in the host [7]. Since the muscle larval stage is the main pathogenic stage of *Trichinella spiralis*, and the antigens excreted and secreted by muscle larvae are easier to prepare, this experiment selected *Trichinella* muscle larvae ES antigen to stimulate the small intestine of mice, with the aim to explore the effect of ES antigen on the host small intestine through the Tuft-IL-25-ILC2 pathway.

There is increasing evidence that luminal worm infection induces epithelial responses, primarily triggered by the activation of tuft cells and the release of IL-25, leading to the activation of type 2 innate lymphocytes (ILC2) and IL-13 secretion [8]. Researchers have reported a positive feedback mechanism, that is, after luminal helminth infection, Tuft cell-derived IL-25 activates lamina propria ILC2, which in turn promotes tuft and goblet cell differentiation in epithelial progenitor cells through IL-13 signaling [9], changing the balance in the small intestine. Two other studies independently defined tuft cells as the primary epithelial sources of IL-25, and highlighted their importance in initiating the type 2 immune response to luminal helminth infection through their interaction with ILC2 [10], which ultimately accelerates parasite clearance. Therefore, in this study, the small intestinal tissue of mice was stimulated with *Trichinella sp*. ES antigen, and an IL-25 blocking group was set up to verify the existence of ES antigen in the Tuft-IL-25-ILC2 pathway. It was found that *Trichinella sp*. ES antigen can activate this pathway to secrete related cytokines. The results of RT-PCR showed that, compared with the control group, the mRNA expression levels of IL-13, IL-25 and IL-25R were increased in the ES antigen stimulation group and decreased in the IL-25 blocking group. Consistent with the above results, it was found that IL-25 might affect the expression of related factors in the Tuft-IL-25-ILC2 pathway in the ES antigen-stimulated mouse small intestine.

Doublecortin-like kinase 1 protein (DCLK1) marks differentiated tuft cells in the small intestine and colon epithelium [11]. Cluster cell-derived IL-25 helps drive the feed-forward Tuft-ILC2 signaling pathway [12], in which ILC2 is activated to produce IL-5 and IL-13, thereby promoting type 2 inflammation. Activation of the Tuft-ILC2 pathway soon leads to tuft cell and goblet cell hyperplasia due to the rapid turnover of the intestinal epithelium. One study reported that *Trichinella spiralis* extract and *Trichinella* ES antigen stimulated enterocytes *in vitro* and demonstrated the increased expression of IL-25 cytokine and elevated number of goblet cells [13]. By stimulating the mouse intestinal Alcian blue-nuclear fast red staining experiment with the *Trichinella* ES antigen, it was also confirmed that the *Trichinella* ES antigen can significantly increase the number of goblet cells in the small intestine. Using immunofluorescence double staining (Dclk1+Pou2f3), we also detected the changes in the number of Tuft cells in the small intestine of mice stimulated with *Trichinella* ES antigen. Compared with the control group, the number of Tuft cells in the small intestine of mice in the *Trichinella* ES antigen-stimulated group increased; in comparison, the number of Tuft cells in the small intestine of mice in the IL-25 blocking group was significantly reduced. Similar to the above, we found that the increased number of goblet cells in mouse small intestine may be related to ES antigen stimulation.

IL-25 is a member of the IL-17 family and is known to be involved in mediating type 2 immunity against gastrointestinal helminth infection [14]. It is one of the activators of intestinal ILC2 cells that can bind to the receptor IL-17RB (IL-25R) on their surface [15]. Thus, ILC2 cells are activated to secrete cytokines such as IL-4, IL-5 and IL-13 to promote Th2 immune response. Therefore, to further explore the role of ILC2 cells in the small intestine of mice stimulated by *Trichinella* ES antigen, we observed the mRNA expression of IL-13, IL-25, RORα, Pou2f3, and IL-25R in the small intestine of mice by RT-PCR. The results showed that, compared with the control group, the mRNA expression levels of IL-13, IL-25, RORα, Pou2f3, and its receptor IL- 25R in the small intestine of ES antigen-stimulated mice were increased; however, the mRNA expression levels of RORα, Pou2f3, IL-25, and its receptor IL-25R in the small intestinal tissue of mice in the IL-25 blocking group were lower than those in the *Trichinella* ES antigen stimulation group. We speculated that the stimulation of IL-13, RORα, Pou2f3, IL-25, and its receptor IL-25R in mouse intestinal tissue by *Trichinella sp*. may be linked to the activation of intestinal ILC2 cells, and IL-25 may stimulate ILC2 cells, prompting them to secrete related cytokines that play an important role in the mouse intestinal tract.

In conclusion, this experimental study found that Tuft cells may be involved in the stimulation of the small intestine of mice by ES antigen of *Trichinella spiralis*, and secreted IL- 25 cytokines to activate intestinal ILC2 cells. Through *in vivo* experiments, it was confirmed that the Tuft-IL-25-ILC2 pathway may exist during the stimulation of the small intestine of mice with *Trichinella sp*. ES antigen. Moreover, we observed that the number of Tuft cells and ILC2 cells decreased after IL-25 was blocked, and the expression of cytokine IL-13 was also decreased, which may provide new ideas for studying the immune mechanism of trichinosis.

## ACKNOWLEDGEMENTS

We thank Professor Lu Yixin from the Heilongjiang Provincial Key Laboratory of Zoogenic zoonosis, College of Animal Medicine, Northeast Agricultural University for his kindly providing *Trichinella spiralis*. The authors wish to thank Dr. Zhang Jinpeng for his technical assistance.

## AUTHOR CONTRIBUTION

HE Ling performed the research and data analysis, Bai Jie contributed to research design and manuscript edition, Napisha·Jureti contributed to Western Blot studies.

